# Membrane Tethering of SepF, a Membrane Anchor for the *Mycobacterium tuberculosis* Z-ring

**DOI:** 10.1101/2022.08.26.505433

**Authors:** Souvik Dey, Huan-Xiang Zhou

## Abstract

Bacterial cell division begins with the formation of the Z-ring via polymerization of FtsZ and the localization of Z-ring beneath the inner membrane through membrane anchors. In *Mycobacterium tuberculosis* (*Mtb*), SepF is one such membrane anchor, but our understanding of the underlying mechanism is very limited. Here we used molecular dynamics simulations to characterize how SepF itself, a water-soluble protein, tethers to acidic membranes that mimic the *Mtb* inner membrane. In addition to an amphipathic helix (residues 1-12) at the N-terminus, membrane binding also occurs through two stretches of positively charged residues (Arg27-Arg 37 and Arg95-Arg107) in the long linker preceding the FtsZ-binding core domain (residues 128-218). The additional interactions via the disordered linker stabilize the membrane tethering of SepF, and keep the core domain of SepF and hence the attached Z-ring close to the membrane. The resulting membrane proximity of the Z-ring in turn enables its interactions with and thus recruitment of two membrane proteins, FtsW and CrgA, at the late stage of cell division.

---

*Mycobacteria tuberculosis* (*Mtb*) is the most infectious bacterium to date, inflicting a quarter of the worldwide population [1]. Most drugs for treating tuberculosis (TB) target cell growth and division. With drug-resistant TB cases increasing rapidly, it becomes even more urgent to better understand the *Mtb* cell division process. As with bacteria in general, *Mtb* cell division begins with the formation of the Z-ring via polymerization of FtsZ and the localization of Z-ring beneath the inner membrane. In the well-studied model bacteria *Escherichia coli* and *Bacillus subtilis* (*Bsu*), two proteins, FtsA and ZipA in *E. coli* and FtsA and SepF in *Bsu*, have overlapping roles as membrane anchors for the Z-ring [2, 3]. ZipA is a transmembrane protein [4], whereas FtsA and SepF are water-soluble proteins with an amphipathic helix for membrane tethering [5, 6]. FtsZ uses its disordered C-tail to bind a structured domain in each of these three proteins for membrane anchoring [6-8]. The role of SepF as a membrane anchor of the Z-ring has also been characterized for the homolog in *Corynebacterium glutamicum* (*Cgl*) [9]. Gola et al. [10] suggested a similar role for the SepF homolog in *Mycobacteria smegmatis* (*Msm*), the latter being often used as a nonpathogenic model for *Mtb*. However, these authors also recognized that the linker between the N-terminal amphipathic helix and the C-terminal FtsZ-binding core domain is much longer in the mycobacterial SepFs (Figures 1a and S1a, b), and may code for additional functions. Here, using molecular dynamics (MD) simulations, we studied the membrane tethering of *Mtb* SepF to delineate its role as a membrane anchor of the Z-ring.

**Figure 1.**
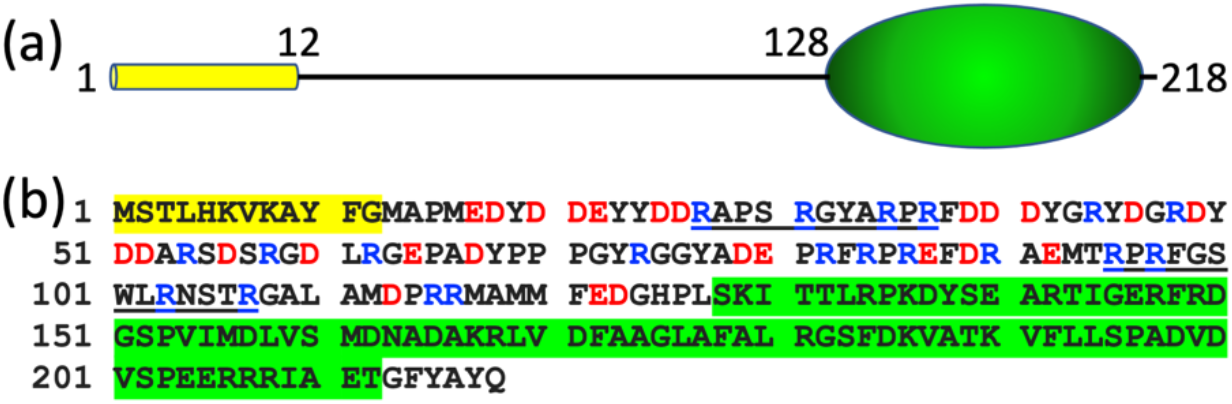
Domain decomposition and sequence of *Mtb* SepF. (a) N-terminal amphipathic helix (yellow cylinder) and FtsZ-binding core domain (green ellipsoid), connected by the linker. (b) Amino-acid sequence, color-coded as in (a). In the linker portion, positively and negatively charged residues are shown in blue and red letters, respectively.

MD simulations have recently become a powerful tool for studying intrinsically disordered proteins (IDPs) in general and their membrane binding in particular. This advance was made possible through GPU acceleration [11] and by selecting a force-field combination [12-14] that is validated for IDPs both in solution and bound to membranes [15-17].

The *Mtb* SepF linker, from residues 13 to 127, is over-represented by charged residues (40%; Figure 1b) and disordered (Figure S2). As a first step to sample the membrane tethering of SepF by MD simulations, we prepared two fragments, encompassing residues 1-50 and 66-124, respectively, near membranes consisting of POPG and POPC at a 7:3 molar ratio, to mimic the highly acidic inner membrane of *Msm* (and presumably also of *Mtb*) [18]. Residues 1-12 were modeled as an amphipathic helix whereas all other residues were disordered. In multiple simulations of SepF1-50 lasting 825 ns each, the amphipathic helix is stably buried at the membrane interface as expected (Figure 2a, b). The nonpolar sidechains of Met1, Leu4, Val7, and Phe11 interact with the acyl chains of the lipids. On the other hand, Tyr10 uses its phenol ring to interact with the acyl chain of one lipid but uses its hydroxyl to form a hydrogen bond with the carbonyl oxygen of another lipid. Moreover, the amines of Lys6 and Lys8 snorkel to the membrane surface to form favorable electrostatic interactions with the headgroups of lipids. The protein chain then climbs out of the membrane. However, from residues 27 to 37, SepF dips back to the membrane surface (Figure 2a). The four Arg residues, Arg27, Arg31, Arg35, and Arg37, in this stretch frequently form electrostatic interactions with the headgroups of POPG lipids (Figure 2c). Tyr33 can also hydrogen bond to lipid headgroups. The Arg residues in the 27-37 stretch are uninterrupted by negatively charged residues; different subsets of these Arg residues tether this stretch to the membrane all the time (Video S1). The membrane contact probabilities of individual residues in SepF1-50 are displayed in Figure 2d.

**Figure 2.**
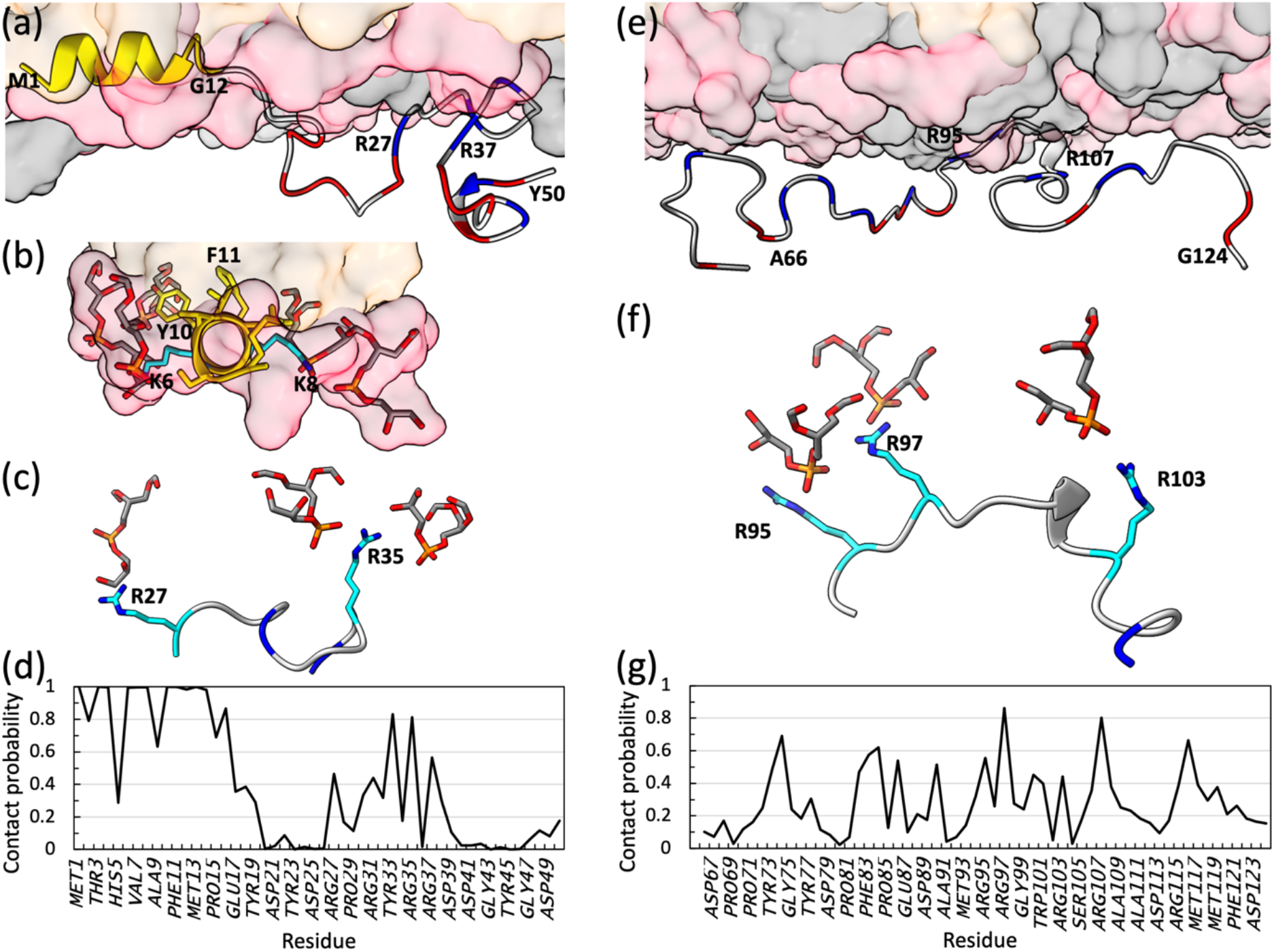
Membrane interactions of two *Mtb* SepF fragments, 1-50 and 66-124. (a) Snapshot showing SepF1-50 bound to a POPG/POPC membrane. Lipids are rendered as surface, with POPG and POPC headgroups in pink and gray, respectively, and acyl chains in orange. For SepF1-50, the first 12 residues are in yellow; the remainder is in gray, except that Asp and Glu are in red and Arg are in blue. (b) View down the axis of the amphipathic helix. Nonpolar residues (including Phe11) take top positions and are buried in the hydrophobic region of the membrane, whereas polar (Tyr10) and charged (Lys6 and Lys8) residues take side position and interact with POPG headgroups (shown as both surface and stick). (c) View into the membrane surface, zoomed on the 27-37 stretch. Arg sidechains interacting with POPG headgroups are shown as stick. (d) Membrane contact probabilities of residues in SepF1-50. A contact was defined as formed when two heavy atoms, one from the protein and one from the membrane, were within 3.5 Å. The contact probability of a residue was calculated as the fraction of saved snapshots in which it formed at least one membrane contact. (e) Snapshot showing SepF66-5124 bound to a POPG/POPC membrane. (f) View into the membrane surface, zoomed on the 95-107 stretch. Arg sidechains interacting with POPG headgroups are shown as stick. (g) Membrane contact probabilities of residues in SepF66-124. The MD simulation protocol was as described in Hicks et al. [17]. Each membrane leaflet contained 154 POPG lipids and 66 POPC lipids; in the SepF1-50 simulations, 6 additional POPC lipids were added to the upper leaflet to achieve parity in area between the two membrane surfaces. The simulation box dimensions were 120 Å ξ 120 Å ξ 140 Å, containing approximately 230,000 atoms. Four replicate simulations were run for SepF1-50, each lasting 825 ns. For SepF66-124, the number of replicate simulation was increased to eight, and the time of each simulation was 1075 ns. After discarding the first 75 ns, snapshots were saved every 20 ps for analysis.

In the simulations of SepF66-124, all the Arg residues frequently form electrostatic interactions with the headgroups of POPG lipids (Figure 2e). This is especially true of the 95-107 stretch, where four Arg residues, at 95, 97, 103, and 107, are uninterrupted by negatively charged residues. This stretch has the closest approach to the membrane surface (Figure 2e), with different subsets of the four Arg residues tethering it to the membrane all the time (Figure 2f; Video S2). The membrane contact probabilities of individual residues in SepF66-124 are displayed in Figure 2g.

The *Cgl* SepF core domain forms a back-to-back dimer, with two symmetric FtsZ-binding sites near the dimer interface [9]. We built a full-length *Mtb* SepF dimer bound to a POPG/POPC membrane, by piecing together the 1-50 and 66-124 fragments taken from the MD simulations presented above and a homology model for the core domain dimer (Figure 3a). In multiple MD simulations of the full-length *Mtb* SepF dimer, the membrane tethering by the amphipathic helices and linkers in both chains is stable (Figure 3b). In particular, the two stretches, 27-37 and 95-107, in the linkers remain membrane-tethered all the time, via Arg residues. The membrane contact probabilities of individual residues in the amphipathic helices and linkers of the full-length dimer are similar to those found in the simulations of the two fragments (Figure S3). A few more residues in the 27-37 stretch simultaneously participate in membrane tethering in the full-length context, possibly because the downstream region is no longer free to move away from the membrane due to the tethering of the 95-107 stretch. The opposite seems to be true for the last few residues of the linker, possibly due to restraints imposed by the downstream core domain dimer. Overall, we find that the membrane binding of the linker, in particular through the 27-37 and 95-107 stretches, reinforces the membrane tethering of SepF afforded by the N-terminal amphipathic helix. The linker of *Msm* SepF also has two similar stretches for membrane tethering (Figure 1c).

**Figure 3.**
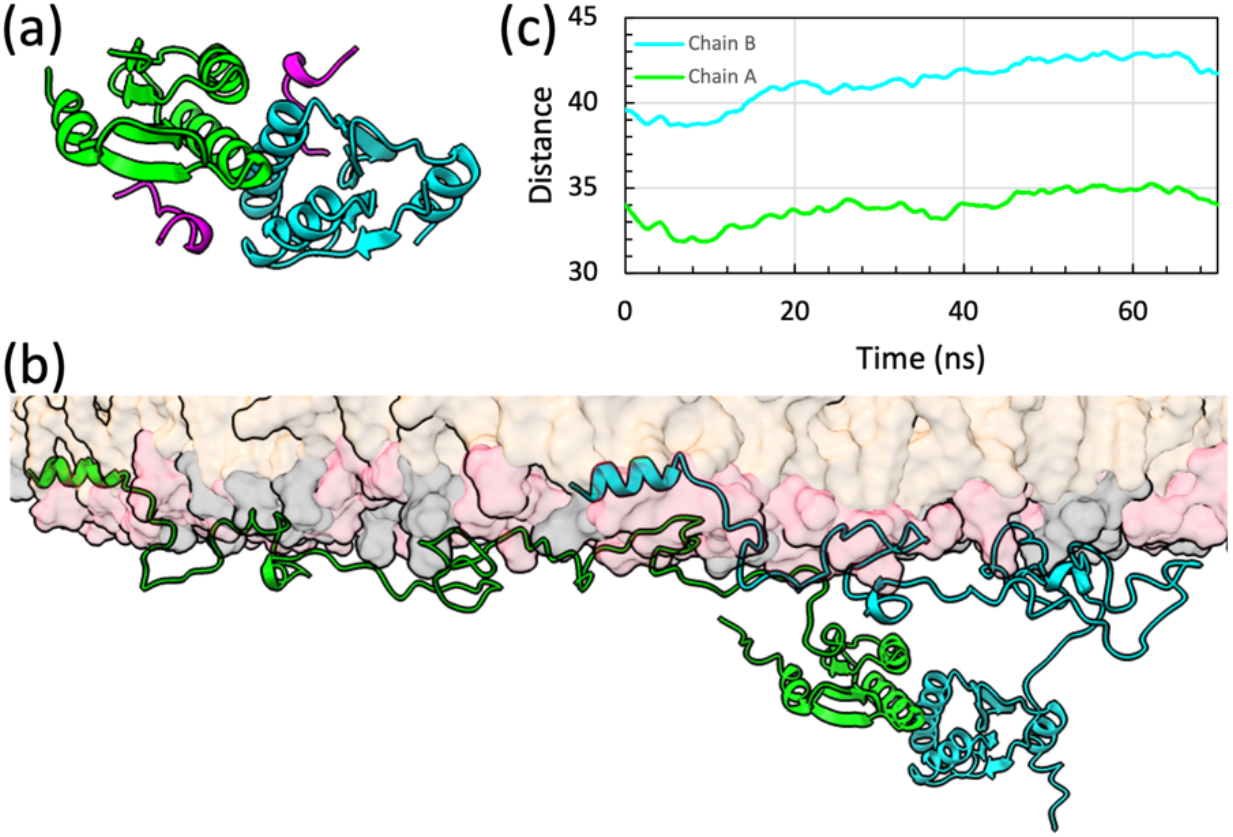
Membrane interactions of full-length *Mtb* SepF dimer. (a) Core domain dimer bound with FtsZ C-tail (in magenta), built by homology modeling using the *Cgl* structure (Protein Data Bank entry 6SAT) [9] as template. (b) Snapshot showing the full-length dimer bound to a POPG/POPC membrane. The two chains of the SepF dimer are shown in green and cyan, respectively. (c) The distances between the centers of mass of the core domains in the two SepF chains and the phosphate plane. The lower leaflet contained 490 POPG lipids and 210 POPC lipids; the upper leaflet had 12 additional POPC lipids. The simulation box dimensions were 214 Å ξ 214 Å ξ 173 Å, containing approximately 930,000 atoms. Four replicate simulations were run, each lasting 70 ns. Snapshots were saved every 20 ps for analysis.

An important consequence of the membrane tethering by the linker is that the core domain is much closer to the membrane surface when compared to the hypothetical situation where the linker is free of membrane tethering. The distances of the core domains from the membrane surface in the MD simulations of the full-length *Mtb* SepF dimer are shown in Figure 3b. They range from 32 to 43 Å. Assuming that Arg107 is tethered to the membrane, there are only 20 disordered residues that act as the spacer between the membrane and the core domain. The mean distance spanned by an *n*-residue disordered spacer can be estimated as

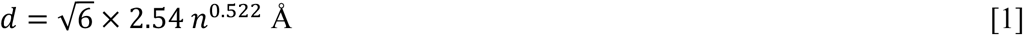

[19]. For our case, *n* = 20 and *d* = 30 Å. Depending on whether the core domain is oriented sideways or directly below the last residue of the linker, approximately 0 to 10 Å should be added, resulting in a range of 30 to 40 Å for the distance between the center of geometry of the core domain and the membrane surface. This estimate agrees well with the simulation results. In comparison, if the entire 115 residues of the linker formed the spacer, the distance from the core domain to the membrane surface would be 74 to 84 Å.

Because the Z-ring attaches to the SepF core domain via the disordered C-tail of FtsZ (Figure 3a) [6, 9], the relatively short distance between the *Mtb* SepF core domain and the membrane surface means that the *Mtb* Z-ring is also close to the membrane. This membrane proximity is necessary because the *Mtb* Z-ring also binds two membrane proteins, FtsW and CrgA (Figure 4), at the late stage of cell division. FtsZ and FtsW interact via their disordered C-tails [20]. The interaction site on FtsZ was identified as a stretch of four consecutive Asp residues, 367-370; the interaction site on FtsW was isolated to a fragment consisting of residues 490 to 524, most likely an Arg-rich stretch from 510 to 516, with sequence RRTRRVR. Because FtsZ binds to both SepF and FtsW via its C-tail, the FtsW Arg-rich stretch must be able to reach a site at the same distance from the membrane as the FtsZ-binding site on the SepF core domain (dashed semicircle in Figure 4). This distance as estimated above is 30 to 40 Å, assuming that the SepF linker is tethered to the membrane as we have found in the MD simulations. The disordered C-tail of FtsW starts around residue 455; the portion of the C-tail from residue 455 to residue 509 (just before the start of the Arg-rich stretch) can easily span a distance of 50 Å (according to equation [1]). If the SepF linker were not tethered to the membrane, the Z-ring would be farther out and the FtsW Arg-rich stretch would be difficult to reach the necessary distance, now increased to 74 to 84 Å.

**Figure 4.**
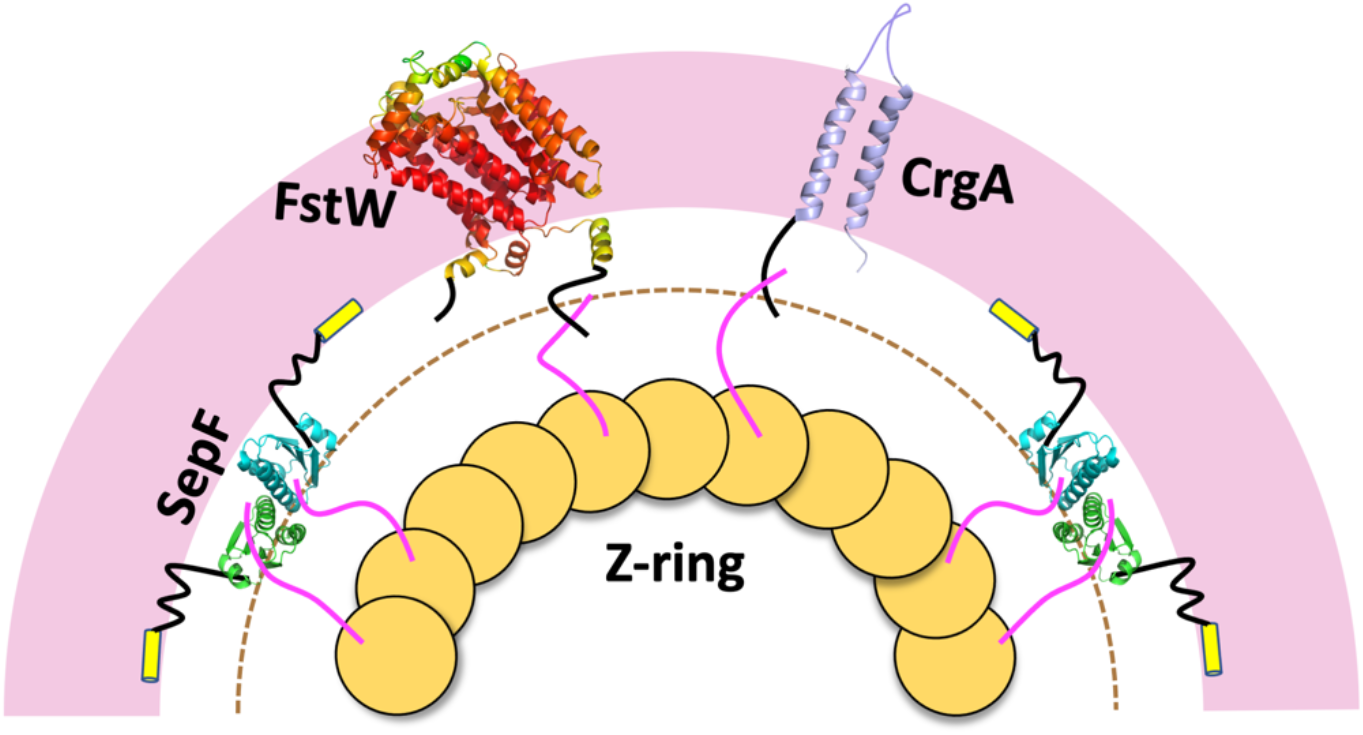
Illustration of the membrane anchoring of the *Mtb* Z-ring by SepF. Membrane proximity of the Z-ring is required by interactions of two membrane proteins, FtsW and CrgA, with the FtsZ. The minimum distance that the C-tail of FtsW and the N-tail of CrgA have to reach in order to interact with the C-tail of FtsZ is indicated by a dashed semicircle. The structure of FtsW is predicted by AlphaFold (https://alphafold.ebi.ac.uk/) [23]; the color spectrum displays confidence levels of prediction, with confidence decreasing from red to yellow to green. The structure of CrgA is from Protein Data Bank entry 2MMU [22].

The FtsZ-binding site on CrgA was isolated to the disordered N-tail [21], which consists of only 29 residues [22]. Given the short length of the CrgA N-tail, the interaction site on FtsZ is likely also the latter’s C-tail, probably the same 367-370 stretch of consecutive Asp residues as found for FtsW interaction. Indeed, the N-tail of CrgA contains a Lys/Arg-rich stretch, from residues 3-9, with sequence KSKVRKK, similar to the Arg-rich stretch found in the FtsW C-tail. There are 20 residues between the Lys/Arg-rich stretch and the first transmembrane helix in CrgA. These disordered residues can easily span a distance of 30 Å, thereby placing the Lys/Arg-rich stretch within the reach of the FtsZ C-tail. However, if the SepF linker were not tethered to the membrane, it would be impossible for the N-tail of CrgA to bind with FtsZ.

In conclusion, our MD simulations have shown that the long linker of *Mtb* SepF reinforces membrane tethering via its Arg residues, especially in the 27-37 and 95-107 stretches. These membrane interactions also effectively shorten the linker, thereby bringing the SepF core domain and the attached Z-ring into close proximity of the membrane. The membrane proximity of the Z-ring in turn enables its interactions with and thus recruitment of two membrane proteins, FtsW and CrgA, at the late stage of cell division.

## Acknowledgements

This work was supported by National Institutes of Health Grants GM118091 and AI119178.

