## Supplementary Figures for "Membrane Tethering of SepF, a Membrane Anchor for the *Mycobacterium tuberculosis* Z-ring"

*Supplementary Material*

(a)

```
Mtb    1   MSTLHKVKAYFG            12
Msm    1   MSTLHKVKAYFG            12
Cgl    1   MSMLKKTKEFFG            12
Bsu    1   MSMKNKLKNFFS            12
           **  :* * :*.
```

(b)

```
Mtb  128   SKITTLRPKDYSEARTIGERFRDGSPVIMDLVSMDNADAKRLVDFAAGLA   177
Msm  124   AKITTLRPKDYSEARTIGERFRDGTPVIMDLVSMDNADAKRLVDFAAGLA   173
Cgl   67   STIVPVELHSFEDAQVIGGAFRDGDAVVFDMSLLSREEARRIVDFAAGLC   116
Bsu   62   SKVVLSEPRVYAEAQEIADHLKNRRAVVVNLQRIQHDQAKRIVDFLSGTV   111
           :.:.  . : : :*: *.  :::   *:.::  :.. :*:*:*** :*

Mtb  178   FALRGSFDKVATKVFLLSPADVDVSPEERRRIAETGFYAYQ    218
Msm  174   FALRGSFDKVATKVFLLSPADVDVTAEERRRIAEAGFYSYR    214
Cgl  117   FALRGKMQKIDSVTFAVVPELSNISTSELERAARIR         152
Bsu  112   YAIGGDIQRIGSDIFLCTPDNVDVSGTISELISEDEHQRW     151
           :*: *.:::: :  *   *   :::    .  :.
```

(c)

```
Mtb   27   R-APSRGYAR-PR           37
Msm   28   RGARAGGYSRRPR           41
           * * : **:* **

Mtb   95   RP---RFGSWLRNSTR   107
Msm   89   RPAPARLGAM-RGSTR   103
           **   *:*:  *.***
```

Figure S1. Sequence alignment of SepF homologs from *Mtb*, *Msm*, *Cgl*, and *Bsu*. (a) The N-terminal amphipathic helix. (b) The FtsZ-binding core domain. Alignment was done using CLUSTAL Omega (version 1.2.4) [24]. Identical residues are indicated by "*"; similar residues are indicated by ":" or ".". The intervening residues between those shown in (a) and (b) make up the linkers. (c) Manual alignment of two Arg-rich stretches in the linkers of *Mtb* and *Msm* SepFs.

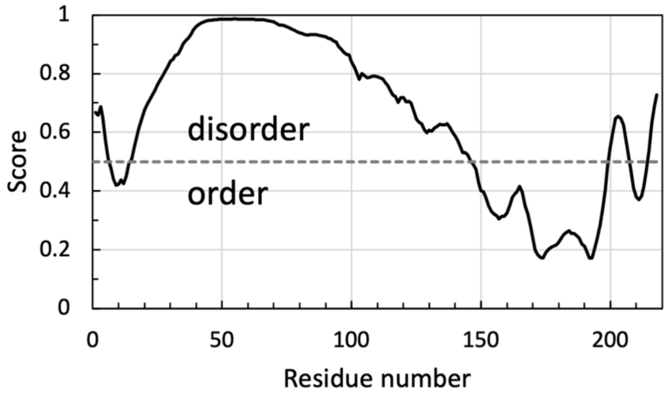


Figure S2. Disorder prediction for *Mtb* SepF using PONDR-VLS2 [25].

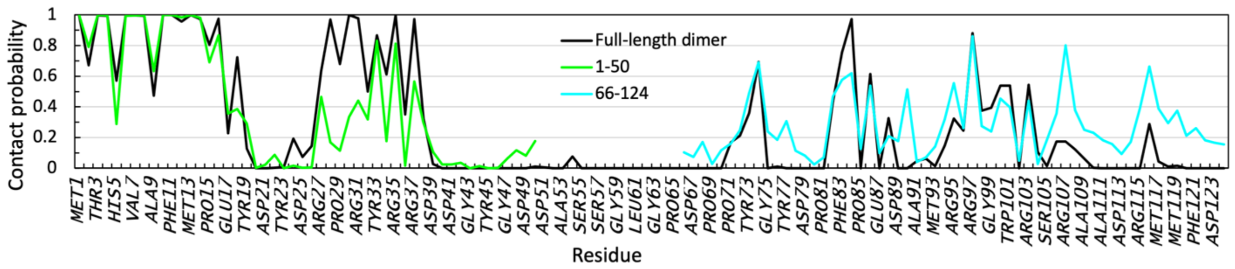


Figure S3. Membrane contact probabilities of residues in full-length *Mtb* SepF dimer, compared with the counterparts in the SepF1-50 and SepF66-124 simulations. The dimer results are the average of the two chains, calculated on snapshots saved after discarding the first 20 ns. The results for the two fragments are the same as those displayed in Figure 2d, g.

Video S1. A 400 ns clip from one of the replicate simulations of SepF1-50 tethered to a 7:3 POPG/POPC membrane. Snapshots were saved every 1 ns and played at 24 per second. The N-terminal amphipathic helix is shown in yellow; the 27-37 stretch is highlighted in green. POPG and POPC headgroups are shown as pink and gray surfaces, respectively; acyl chains are not displayed. Arg and Lys sidechains are shown as blue sticks when any of their heavy atoms are within 3.5 Å of any lipid heavy atoms; POPG (or POPC) headgroups are shown as pink (or gray) sticks when any of their heavy atoms are within 3.5 Å of any Arg or Lys heavy atom.

Video S2. Corresponding video for SepF66-124 tethered to a 7:3 POPG/POPC membrane. The 95-107 stretch is highlighted in green.
